# Digital data storage on DNA tape using CRISPR base editors

**DOI:** 10.1101/2023.02.07.527074

**Authors:** Afsaneh Sadremomtaz, Robert F. Glass, Jorge Eduardo Guerrero, Dennis R. LaJeunesse, Eric A. Josephs, Reza Zadegan

## Abstract

While the archival digital memory industry approaches its physical limits, the demand is significantly increasing, therefore alternatives emerge. Recent efforts have demonstrated DNA’s enormous potential as a digital storage medium with superior information durability, capacity, and energy consumption. However, the majority of the proposed systems require on-demand *de-novo* DNA synthesis techniques that produce a large amount of toxic waste and therefore are not industrially scalable and environmentally friendly. Inspired by the architecture of semiconductor memory devices and recent developments in gene editing, we created a molecular digital data storage system called “DNA Mutational Overwriting Storage” (DMOS) that stores information by leveraging combinatorial, addressable, orthogonal, and independent *in vitro* CRISPR base-editing reactions to write data on a blank pool of greenly synthesized DNA tapes. As a proof of concept, we wrote both a bitmap representation of our school’s logo and the title of this study on the DNA tapes, and accurately recovered the stored data.

---

As digital information production grows exponentially and the industry approaches physical limits, high density long-term storage solutions are necessary.^1,2^ DNA as an alternative to the archival storage medium offers several potential advantages, including higher density and retention, and lower energy consumption compared to the state-of-the-art memory materials.^3-6^ *De novo* DNA synthesis has enabled the development of DNA-based memory technologies^7-10^ and the 2018 Semiconductor Synthetic Biology (SemiSynBio) Roadmap predicts that the speed and cost of DNA synthesis will improve dramatically in the future. Additionally, automation of writing and reading processes may improve the portability and scalability of DNA memory. However, *de-novo* DNA synthesis methods are unlikely to meet the scalability required for high-volume memory manufacturing, and generate massive amounts of toxic waste, which is not sustainabile.^11-16^ For context, our calculations demonstrated that storing 5 minutes of a 1080p YouTube video stream in commercially acquired DNA costs over 7 million dollars, consumes over 100KWh of energy, takes over 4 days, and produces over 15 liters of toxic waste (Supplementary S1). This implies, to store every bit of information generated by 2030 (∼6e+23 Bytes) in synthetic DNA, the current technology would produce nearly 85 petaliters of hazardous waste, which surpasses the volume of water that the Mississippi River empties into the Gulf of Mexico over 40 years.^17^ While the majority of academic and industrial efforts target scalability, the DNA memory community has mostly ignored the disruptive environmental consequences of large-scale *de-novo* DNA synthesis. For the DNA-based memory innovations to become mainstream the environmental concerns need to be addressed.

Our molecular data storage system, that we coined ‘DNA Mutational Overwriting Storage’ (DMOS), uses combinatorial, addressable, orthogonal, and independent in-vitro CRISPR base-editing reactions to overwrite data on the pre-defined domains of existing DNA. Akin to conventional magnetic tape architecture, the ‘blank DNA tape’ consists of multiple DMOS registers and each register contains a set of 16 domains (bits) –including ‘state’ and ‘index’ sections. We employ CRISPR base editing^18^ to write the information by mutating the domain state from the unmutated state (0) to the mutated state (1). Using nanopore sequencing, we recover the ‘mutational signature’ of each DNA tape register that informs our error-correction and coding schemes to retain the data accurately and precisely. As a proof of concept, we wrote 1250 bits of data, including both a bitmap representation of our school’s logo and the title of this paper on multiple DNA registers, and recovered the stored data with 100% accuracy. This work is the first demonstration of writing digital data in the form of sequence edits at precise locations of a pre-existing blank pool (called DNA tape) of greenly synthesized DNA registers.

## The writing mechanism

To write the data on DMOS bits, we developed a programmable molecular writer system that uses CRISPR base-editing reaction. During the base-editing reaction, the CRISPR effector Cas9 first recognizes and binds to a 3 bp sequence known as the protospacer adjacent motif (PAM) and a 20 bp-long sequence complementary to the targeting guide RNA (gRNA). The 20bp gRNA complementary sequences within the double-stranded DNA registers make up the ‘state’ sections of each domain (Supplementary S2, Fig. 1A, S1). The resulting complex forms a nucleotide structure known as an R-loop. During CRISPR-Cas9 gene editing, the formation of the R-loop would trigger the Cas9 to generate a double-strand break in the DNA molecule. However, in a modified base-editing reaction we use a mutant form of Cas9 known as dead Cas9 (dCas9) recognizes and binds to specific DNA targets to form an R-loop while keeping the DNA intact^18^ (Supplementary S3, Fig. S2). We then used a mutagenic protein APOBEC3A to modify the domain sequences at highly susceptible displaced DNA strands of registers independently in the select positions. These reactions mutate deoxycytidine (dC) to deoxyuracil (dU), which is subsequently converted permanently to deoxythymine (dT) in the target domains with a resolving biochemical reaction (Supplementary S2, Fig. 1A). We designed and experimentally validated a set of 16 unique state sequences where i) Cas9 exhibited robust double-strand cleavage activity at the target location and allowed each gRNA to form a stable R-loop with the target state sequence^19^; ii) each displaced DNA sequence contained at least two dTdCdR nucleotide motifs^20^ that were recognized by APOBEC3A (where dR is a dG or dA nucleotide); and iii) those motifs located in a mutagenic hot-spot and 6 nt away from the PAM region. The last criterion is a result of the dCs located close to the PAM (within ∼6 bp) having significantly lower mutation rates (Fig. S3). To write the data, the dCs across the state section of a DMOS bit are mutated to dTs if and only if Cas9 with the gRNA targeting the state segment of that specific bit is included in the reaction (Fig.1). To read the state of the bit we determined whether dCs in that bit were mutated (1) or unmutated (0). APOBEC3A efficiently mutates dCs in single-stranded DNA or R-loops, therefore only the dCs in registers with a bound dCas9 will be mutated, while those in registers without dCas9s will remain unmutated. Indeed, analysis of targeted versus non-targeted bit sequences confirmed that APOBEC3A only affected DNA when bound by dCas9. The other state segments remain base-paired, and therefore the mutations happen only in the displaced strand of an R-loop of a targeted bit (Fig. S1).

**Figure. 1:**
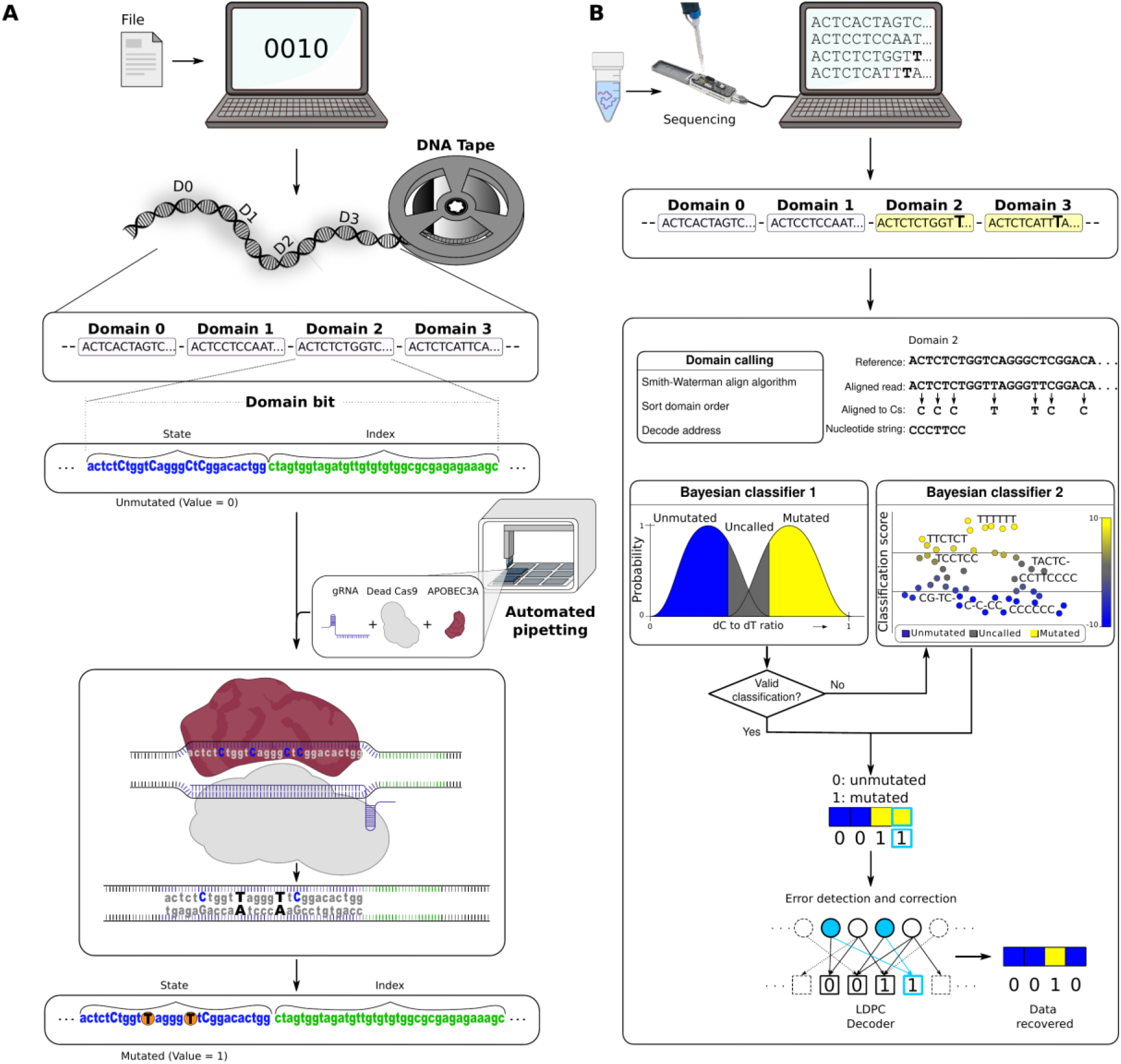
Schematic view of writing data onto DMOS DNA tape. To write and read the data the encoder uses binary files and converts them into byte arrays with codewords. The coding protocol informs the writer system and determines the desired edits on the state (bit) sections of the DNA registers resulting in changing the states of bits from 0 to 1 on DNA tape. A) Writing data on blank magnetic tape. Input data is converted to the binary message and inform the mutation process. DMOS uses a programmable molecular writer system that drives CRISPR base-editing reactions. CRISPR/dead Cas9 (dCas9) accompanied by APOBEC3A drives mutation reactions for the parallel re-writing of data in state sections of DNA tape registers. B) To read the data, we use nanopore DNA sequencing. The output of sequencing reads is decoded by first performing a local alignment (Smith-Waterman alignment) followed by a Bayesian analysis to determine the mutated states and convert the data back to the binary form.

## The architecture of DMOS blank tape registers

Like writing data onto blank tapes, the DMOS overwrites the sequence (state) of the DNA domains to write the data. DMOS tape contains several registers, and each register consists of 16 permuted predetermined domain bits (Fig.1A, S4A). Each domain bit consists of a 23 bp-long ‘state’ section (PAM and sequence recognized by one of the gRNAs) and a unique 40 bp ‘index’ sequence (Figure. 1A). This architecture spaces out the bits far enough for compatibility with the writer system, resulting in independent interactions and reduced crosstalk.

We used the indexes for addressing, increasing error-tolerance, and localizing a bit along a register during sequencing (Fig. 1A). We tested three addressing methods (Supplemental S4) for data recognition in the same sequencing run, allowing to generate a DMOS block with highly differentiable DMOS registers in one single sequencing data pool and reduce the computation time and costs (Figure.S1A). For the first method, we used barcode-based addressing scheme that exploited the 36 combinations of 6 unique barcodes to recognize the registers in the pool (Fig. S4B). However, this reduces the system scalability and limits recognition of registers. Therefore, our second method uses a coding scheme (Supplementary S4) for shuffling (permutating) domain positions in each register (Fig. S4B) that helped to determine the identity and order of the bits in each individual register (hereby called domain-calling). To construct the permutations, we used two shuffling schemes; Low-entropy addressing scheme and High-entropy addressing scheme.^21^ The Low-entropy addressing scheme is a lexicographic representation of the permutation order where we kept the first 10 domain positions constant and shuffled the last six domains. The High-entropy addressing scheme was designed based on permutating the order of domain positions in the entire register (Fig. S4B). The low-entropy addressing scheme leads to inefficient sequence analysis (see Fig. S4B). Therefore, we used the High-entropy addressing scheme that permutates the order of bit positions in the entire register (Fig. S4B). This scheme ensures efficient bit recovery from the sequencing data in our experiments, albeit we assume when used to store big data this scheme may encounter similar challenges as the low-entropy addressing scheme. To increase the data density per sequencing run, we created a DMOS block (tape) of 48 registers (Fig. S4B). Finally, we used bacteria to synthesize the DNA registers used for DMOS.

## The orthogonality and modularity of the DMOS writer system

To parallelize data writing, we used a modular DMOS writer system to independently transduce the digital data to the corresponding mutational domains across multiple registers of the DNA tape (Fig. S5A,S5B). We screened the cross-reactivity of the writer system and confirmed that the mutation reactions at the desired locations were independent (Fig. S5A). We ran the experiments based on a programmed targeting of individual DMOS bits as a function of the dC to dT mutation rates across the 16 distinct bits on a single register. As shown in Fig. S5A, we confirmed that multiplexed targeting of neighboring bits did not change the mutational signatures compared to bits reacted one-at-a-time (Fig.S3B,S5A,S5B,S6). These values also informed our algorithm and convert nanopore sequencing reads to digital data based on ‘called-bits’ to represent the presence or absence of the mutational signature (Fig. S1, S5, S6). We used a Bayesian classifier to analyze the sequencing results (reads) and generated the mutational signature of each bit (Fig. S6). Observed bit mutation rates that were significantly higher or lower in comparisons to the thresholds determined by the mutational signatures (Table S3) represent 1 or 0, respectively, while those near the threshold were called uncertain (Fig. S5, S6) and subjected to more additional and more stringent Bayesian analysis of their read sequences to infer their state based on that bit’s mutational signature. Overall, our results highlighted orthogonality, independency, and reproducibility across bits of DMOS registers (Supplementary S5).

We created a bitmap representation of our school’s name (Joint School of Nanoscience and Nanoengineering) in 512 pixels across 32 DMOS registers (Fig. 2). We plotted out the desired locations of the 0s and 1s and used an open-source Opentrons robot to automate the delivery of the gRNA/dCas9 complexes accordingly. Figure S7 shows the normalized mutation probability graph versus the dC-to-dT mutation rate for individual DMOS bits. We ran the two rounds of Bayesian classifiers to differentiate Mutated and Unmutated states for every single sequencing read in each DMOS register (Fig.S7). Next, we assigned digital states to the corresponding register bits to increase bits calling accuracy for 100 nanopore sequencing intervals (Fig. 2). After 20k reads, we could partially recover the intended bitmap. Notably, the ‘bit recovery’ curve reaches a plateau after 10k reads and additional reads result in an insignificant increase in data recovery (Fig. 2 and supplementary S6).

**Figure. 2:**
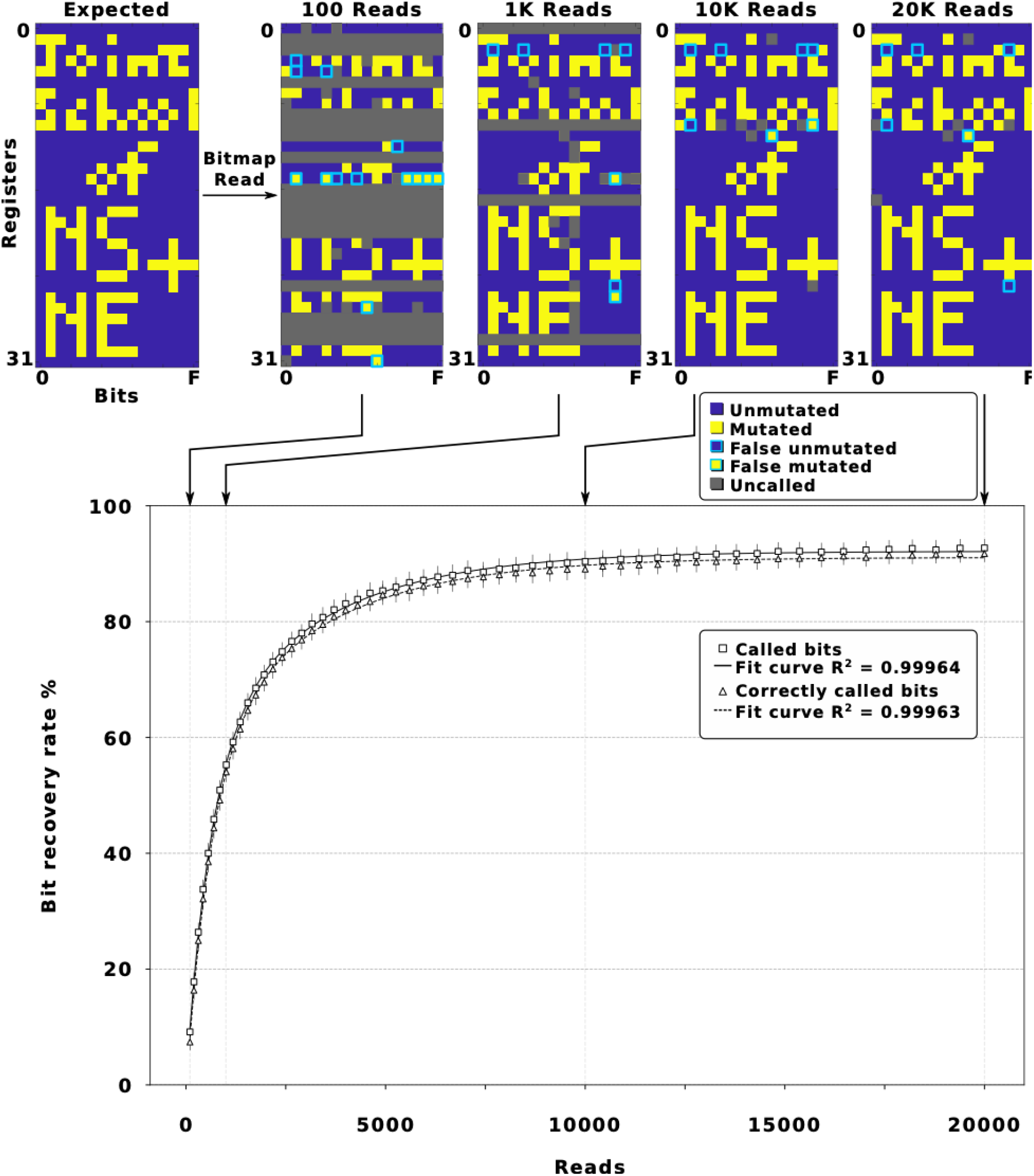
Bitmap representation of the logo of our school (the Joint School of Nanoscience (NS) and Nanoengineering (NE)) written on DMOS tape. The DMOS decoder records snapshots in every 100 nanopore sequencing read. Here, an example of these snapshots at 100, 1000, 10,000, and 20,000 reads is presented. The bootstrap analysis was performed with 250 replicate selections from different positions, which is represented as the recovery rate of called bits and correctly called bits vs. reads.

Additionally, we assessed the robustness of the method via bootstrap analysis of 250 randomly selected sequencing data streams, where the data streams (containing a fixed number of reads) were randomly pulled from the sequencing data output that contained the full 20k reads. The generated average called-bits values show the same behavior around 10k reads with a standard deviation of 91.37±1.7 percent (Fig.2). Therefore, we assume the continuation of sequencing to achieve 100 percent data recovery will be computationally costly, if possible (Fig.2). This finding justified the utilization of error-correction codes to achieve acceptable data recovery rate for memory purposes.

## The tradeoff between writing overheads and reading recovery rate

Informed by previous reports^22^ and our statistical bootstrapping analysis, and to reduce the overhead costs of the system, we studied the tradeoffs of the writing overhead and reading recovery rate. The Writing overhead is the amount of added data redundancy that ensures data recovery during the reading process. We employed coding strategies that account for domain-specific coding maps and error correction codes while maximize the bit capacity of the registers to match the domain length of the proposed architecture. To improve data reliability, we used low-density parity-check (LDPC) codes that set the error threshold close to theoretical Shannon capacity limits. The code includes a pre-processing step that adds LDPC error correction to the digital binary contents. Leveraging our bitmap representation sequencing data, we ran an error-recovery simulation for different LDPC codes and generated the models of the error recovery rate for each decoder.^22^ We generated 1000 random data pools and evaluated the studied error-recovery models of each decoder (Fig. S8). Then, we created a library of Protograph and Regular LDPC codes that we used in the subsequent studies (Supplementary S7) (Fig. S9A, S9B). Fig. S8 represents the required error correction overhead for the maximum data recovery rate per errors. These findings suggest the more the error-correction overhead, the faster, the less costly, and the more accurate the data recovery (Supplementary S8, Fig. S8, S9A, S9B, S10). However, it is notable that the block’s storage capacity and hence the cost of wet-lab experiments is affected by increasing redundancy (Figure. S5A). Therefore, we concluded that adding 25 percent redundancy results in reconstructing the data while keeps the overhead at minimum (Fig. S8, S9A).

## Writing and reading digital information on DNA tape

We wrote the title (in ASCII) of this paper ‘Digital data storage on DNA tape using CRISPR base editors’ in the form of sequence edits in one DMOS tape block containing 48 registers (Fig.3). We developed a semi-automated coding platform (herein called D_coder) that utilizes the Protograph LDPC code with 25 percent redundancy. Next, the D_coder generated a file with the list and address of required mutations and communicated with the Opentrons robot to conduct the targeted base editing reactions on the DMOS registers (Fig. 3, S6). After the writing was done, we used nanopore sequencing to read the DNA pool sequences and retrieved the binary contents.

**Figure. 3.**
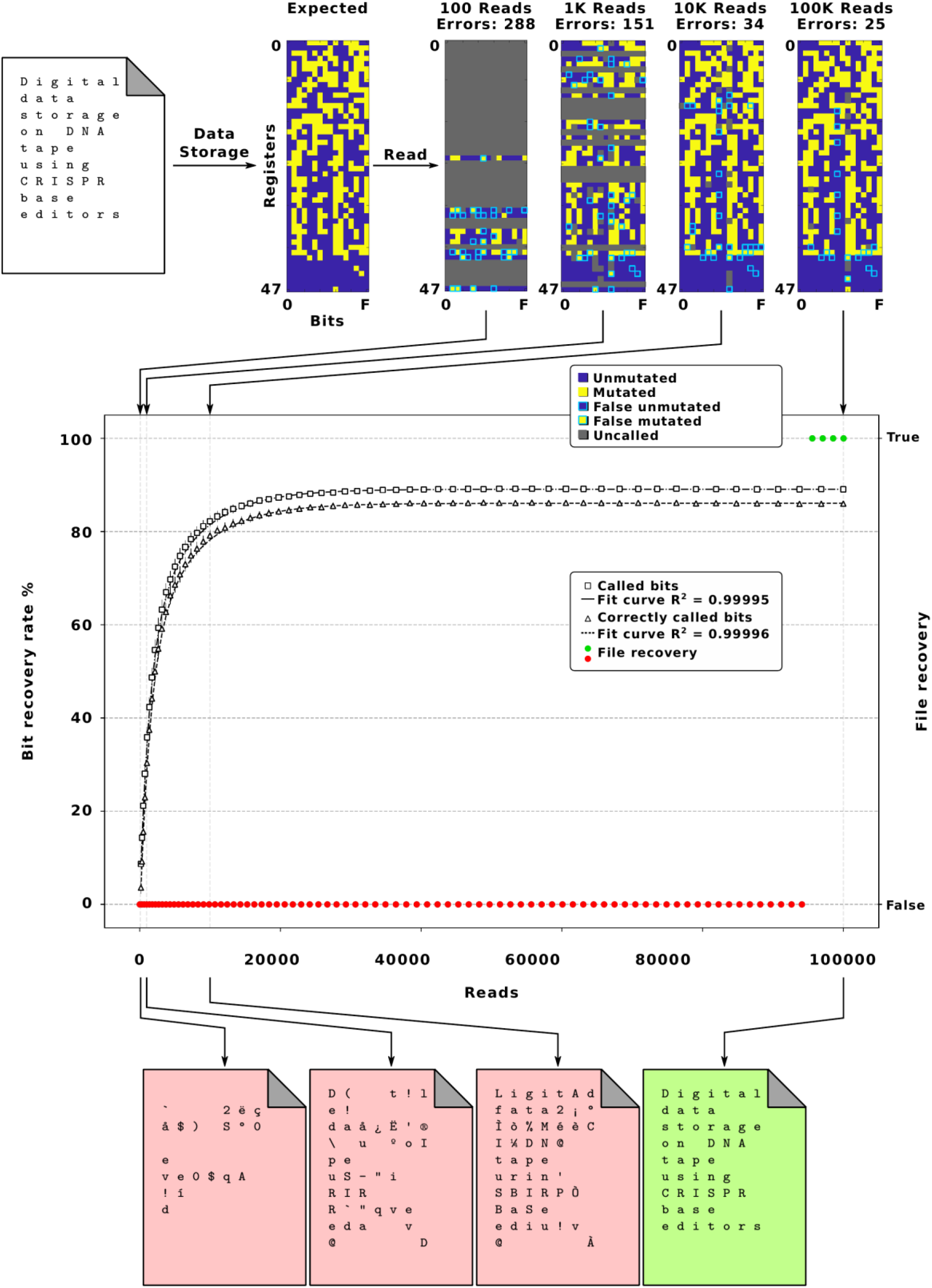
Writing the title of this study on DMOS tape. We converted the title of the paper from ASCII to binary, added error-correction, performed the mutation experiments, sequenced the DNA blocks, and recovered the data. DMOS decoder captured the snapshots of sequencing data streams every 100 intervals. The bootstrap analysis was performed with 250 random replicates of randomly selected DNA sequencing data fragments from the while data. The graph represents the DMOS recovery rate for called bits, correctly-called-bits and file recovery vs. reads.

We evaluated and normalized the probability of the mutations for each isolated DMOS bit and identified their precise mutational signature profiles (Fig. S11). We recorded the results in real-time and converted the sequences to the binary data, translated it to a text file during 100 read intervals, and evaluated the recovery of the file. Like previous study, the bootstrapping of called-bits led to insignificant improvement in data recovery after ∼22K reads (SD 87.37 ± 0.62 (%). Based on our calculations, we expected that our decoder will be able to recover the file when the data steam includes 20k-100k reads. Therefore, we stopped our sequencing data acquisition at 100K reads and using the D_coder fully restored the original file with 100% accuracy and completeness (Fig. 3).

Finally, we ran multiple simulations to assess the performance of several Protograph and Regular LDPC codes with different redundancy percentages that will inform future studies (Supplementary S8, Fig. S8 and S9). E.g., by using Protograph LDPC error-correction with redundancies of 33 and 50 percent, the decoder would be able to fully recover the stored data after 50K and 3K reads, respectively (Figs. S8, S9C).

## Conclusion

‘DNA Mutational Overwriting Storage’ (DMOS) tape leverages the architecture of semiconductor memory devices and recent developments in gene editing technologies to write digital data in form of precise DNA sequence edits on pre-existing DNA strands. We synthesized DNA tape registers in bacteria en-masse, therefore DMOS has the potential to bypasses some challenges of conventional DNA based data storage including generation of toxic wastes, scalability, editability, and rewritability.^23-30^ E.g., in most systems DNA is chemically synthesized, user needs access to DNA synthesizer machines in order to write a new data, and editing of the recorded data requires the synthesis of the whole or significant portions of the DNA pools. Since majority of these challenges are due to the need for de-novo DNA synthesis, a few efforts have focused on alternative DNA based storage media.^31-35^ Some recent studies focused on information stored in DNA nanostructures with rewriting capabilities.^36-38^ E.g. 56 bits of data have been stored using hairpins and overhangs on a long single-stranded DNA genome of M13 while reported retention of 90 percent.^39^ In another report, 20 bytes (160 bits) of digital data was stored in DNA nanostructures and read back using super-resolution microscopy with 100 percent data retention.^40^ Another study stored 14KB of data in form of DNA backbone nicks of pre-existing DNA hard drives.^13, 30^ While promising, majority of these effort lack scalability or sustainability.

The registers that constitute the DMOS DNA tape are universal, and therefore to write new data, DNA synthesis is not required. Additionally, since the registers are amplified in bacteria or via PCR, the reactions do not generate large amount of toxic waste.^3,7-10,13,31-36^ DMOS uses biological enzymes that are inherently slow when compared to semiconductors and hence latency is an issue that needs to be addressed in future studies. Additionally, DMOS sacrifices data density for sustainability and error-tolerance as individual nucleotide mutations are unlikely to affect the data integrity. However, by expanding the gRNA library or multiplexing, DMOS may theoretically be scaled to high capacity.^11-16^

## Methods

### sgRNA synthesis

The coding oligos were ordered from Integrated DNA Technologies. These oligos were used for generating the 16 gRNAs that target the modular domain state sequences. Each sgRNA was synthesized using the S. pyogenes EnGen© sgRNA Synthesis kit (NEB) with 1 uM concentrations of oligo bits. The synthesized RNA was purified using RNAClean XP magnetic particles (Beckman Coulter), and concentrations were recorded using a Nanodrop Lite Spectrophotometer (Thermo scientific).

### Enzymatic writer protocol

We performed the deamination reactions by dispensing 1.5 uL of RNP and 1.5 uL of the DNA template in a 15.5 uL reaction with Cas9 buffer [200nM HEPES, 1M NaCl, 50 mM MgCl2, 1 mM EDTA, pH 7.4]. We prepared the master mix of mutation reaction following this condition: 8.5 uL of nuclease-free H2O, 1.5 uL of 10x Cas9 buffer, 1 uL (40 units) of RNAse Inhibitor Murine, and 0.5 uL of BSA. The APOBEC3A and BSA materials used were purchased from the NEBNext© Enzymatic Methyl-seq Conversion Module kit from New England Biolabs (NEB). The reaction volume was then scaled down to 15.5 uL and modified from the original deamination protocol from the kit. The reaction mix was centrifuged and incubated inside a MiniAmp thermocycler at 37 °C for 3 hours per the standard protocol.

To stop the reaction, we treated with proteinase k following this condition: 1 uL (0.8 units) of proteinase k and incubated at 56 °C for 10 minutes. The samples were purified using AMPure XP magnetic beads following standard protocols. After eluting the sample from the beads, the DNA was treated with Lambda exonuclease to degrade the unmutated strand. This reaction was performed following this condition: 1 uL of Lambda Exonuclease and 5 uL of commercial 10x Lambda Exonuclease buffer for a total reaction volume of 50 uL with nuclease free H2O. The reaction was performed at 37 °C for 30 minutes and heat-inactivated at 75 °C for 15 minutes. Accordingly, the samples were purified using AMPure XP magnetic beads. The mutated samples were addressed to amplification using primers and Q5U from NEB following the PCR setting of annealing temperature of 63 °C and an extension of 72 °C at 45 seconds. The schematic view of this protocol is depicted in Figure.S3. The products were purified, thoroughly mixed in a femtomolar scale, and sequenced through nanopore sequencing.

### Sequencing run and basecalling

Sequencing was performed using an Oxford Nanopore MinION Mk1B nanopore sequencer supported with the MinKNOW software. We prepared a library of DMOS registers with stored files using the ligation sequencing kit “LSK-110” and the sequencing run was carried out on R9.4.1MinION Flongle flow cells from Oxford Nanopore Technologies at default settings on MinKNOW. The fast5 raw signal files were basecalled using Guppy basecaller 6.1 for high-accuracy basecalling on a laptop with Alienware m15 R4 1TB SSD with an Intel i7 10750H CPU, 16 GB of RAM and dedicated NVIDIA GeForce RTX 3060 GPU in the super high accuracy (sup) mode. The generated fastq files were binned into pass or fail folders based on their q-scores. Only the reads that have passed the q-score threshold were analyzed.

### Orthogonality tests on DMOS template

We tested the orthogonality of the DMOS writer system following these conditions: (1) Including dCas9, APOBEC3A, and gRNAs that targeted all 16 DMOS bits simultaneously across the register (triplicate); (ii) no dCas9, no APOBEC3A (triplicate); (iii) including dCas9, APOBEC3A, and gRNAs targeting individual DMOS bits per the DMOS register (triplicate). The mutation reaction performed for all experimental and control reactions under the same standard mutation condition. Next, the reaction was stopped individually with 1 uL of proteinase k (0.8 units) and purified using AMPure XP magnetic beads accordingly. The purified samples were treated with Lambda exonuclease and purified using the standard protocol. Each reaction was amplified and purified individually, and an equimolar from each sample was taken and combined together for the nanopore sequencing. Barcoded primers were used for addressing of DMOS register in this experiment and listed in Table S1.

### Software development

We developed our DMOS D-coder using Python language and the Spyder IDE. The error-correcting layer uses the Protograph LDPC library (https://github.com/shubhamchandak94/ProtographLDPC).^42-45^ To design our LDPC code, we selected the Protograph type accumulative repeat by 4 jagged accumulate to define the Generator and Parity Check matrices, with a message-code ratio of 3/4, expansion factor 96. These parameters constructed an LDPC code that uses 576 bits per message (72 byte) and 768 bits per codeword (96 bytes). We developed a Python script to communicate with the LDPC library that allows the conversion of the intermediate binary files for input/output and capture the diagnostic signals of the LDPC decoder.

The DMOS software layer uses two main modules to retrieve the binary file: DMOS decoder and LDPC decoder. The DMOS decoder was written in C++ using the QtCreator IDE, and uses the Smith-Waterman algorithm (https://github.com/mbreese/swalign/) to align DNA sequences.^44^ We list all the threshold values used in the first Bayesian step (Table S3), and the second Bayesian step uses trained data available in the code repository. We created a graphical user interface using PyQt5 to easily select the input samples and configure the parameters for the DMOS decoder. We developed the simulation scripts in Python language and used standard libraries such as Numpy, Matplotlib, and statistics.

### Automation of writing via OT-2 pipetting robot

We used the Opentrons OT-2 pipetting robots for the automated data writing procedure into DMOS tape. This procedure requires the following plate preparations: We reserved one plate for the dCas9 library of the 16 mutational bits, the second plate contains individual blank DNA registers with addresses; located in separate wells, and the last plate includes a rest master mix content. We developed Python scripts for customizing the mutational list file map and delivering it to the following steps: First, the robot locates the target registers in separate pools. Next, it takes volumes of 1.5uL from the dCas9 library plate and mixes them in the master mix plate. This step is followed by taking 1uL from the master mix and depositing it into the selected register pool. Finally, we incubated the reaction at 37°C for 1 hour at the thermocycler. The samples were addressed for the clean-up step in which we used multichannel tip robots to deposit 30uL of AMPure XP beads into our mixture before activating a magnetic rack for 2 minutes. Next, the magnetic rack was engaged, the supernatant discarded, and the beads were eluted with an elution buffer. The pure DMOS registers pool moves forward with nanopore sequencing.

### Encoding of a file onto DMOS register using DMOS writer system

Following the predetermined map of the mutation list to write the data on registers (Figure.6), the dCas9 RNP pool was prepared and using the Opentrons pipetting robot distributed on the corresponding DMOS registers. Each RNP mixture has a final concentration of 50 nM to preserve the final concentration of having 10 times concentrated RNP to register in every single reaction. Next, the APOBEC3A was added to each reaction and incubated for 3 hours at 37°C inside a veriflex Thermocycler. The reactions were stopped using 1 uL of Proteinase K to degrade the dCas9, followed by individually purifying samples over AMPure XP beads. The purified samples were each treated with Lambda Exonuclease and purified. The purified samples amplified following this PCR setting; primary denaturation 98°C for 30 seconds, denaturation 98 °C for 10 seconds, annealing 63°C for 20 seconds and an extension 72 °C for 45 seconds for 30 cycles followed by final extension at 72 °C for 2 minutes. The registers were purified using standard AMPure XP bead protocols. The samples were combined with an equimolar concentration from each register and sequenced using a nanopore sequencing.

## Acknowledgments

We thank our colleagues Shyam Aravamudhan, Daniel Herr, and Micheal Reed for their support and suggestions. This work is funded through NSF SemiSynBio-II: DNA Mutational Overwriting Storage (DMOS) (award # MCB 2027738) and in part supported through NIH award # R35GM133483. This work was performed at the Joint School of Nanoscience and Nanoengineering, a member of the Southeastern Nanotechnology Infrastructure Corridor (SENIC) and National Nanotechnology Coordinated Infrastructure (NNCI), which is supported by the National Science Foundation (award # ECCS-1542174).

## Data and code availability

The code and data are available at: https://github.com/SBMI-LAB/DMOSEncoder, https://github.com/SBMI-LAB/DMOSDecoder, and https://github.com/SBMI-LAB/DMOS_data.

## Notes

### Competing Interest Statement

The authors have declared no competing interest.

