## Supplemental File for "Digital data storage on DNA tape using CRISPR base editors"

nature materials

### S0. General materials

The DNA barcode primers, 5' phosphate modified reverse primer, 5' phosphorothioate-bond modified forward primer, and gRNA synthesis templates were purchased from Integrated DNA Technologies (IDT; Coralville, IA) and received as dry pellets. The primer oligos were dissolved in nuclease-free H2O up to a concentration of 100 uM and stored at -20 C for future use. The DMOS blank tape and the registers used for the school logo and title encoding experiments were purchased from TWIST Bioscience (San Francisco, CA) as dry pellets, resuspended in 0.1X TE buffer from Thermo Fisher Scientific (Waltham, Massachusetts) and stored at -20 C.

The FastDigest EcoRI restriction enzyme (catalog number FD0274) and FastDigest Bsu15I restriction enzyme (catalog number FD0144) were purchased from Thermo Fisher Scientific (Waltham, Massachusetts), and the PBR322 plasmid (catalog# N3033S) was purchased from New England Biolabs (Ipswich, MA) and stored at -20 C. The high-efficiency NEB 5-alpha Competent E. coli cells (catalog# C2987H) were purchased from New England Biolabs (Ipswich, MA) and stored in -80 C. Cloning results were confirmed via Sanger Sequencing by Azenta (South Plainfield, NJ).

The dCas9 (catalog number 1081058) and Cas9 proteins (catalog number 1081066) were purchased from Integrated DNA Technologies (IDT; Coralville, IA). The enGen sgRNA synthesis kit (catalog number E3322), lambda exonuclease (catalog number M0262S), and Q5U Polymerase master mix(catalog number M0597) were all purchased from New England Biolabs (NEB, Ipswich, MA). The APOBEC3A enzyme and BSA buffer were extracted from the NEBNext Enzymatic Methyl-seq Conversion Module (catalog number E7125) purchased from New England Biolabs (NEB; Ipswich, MA). The AMPure XP magnetic particles (catalog number A63881) were purchased in 60 mL volumes from Beckman Coulter (Indianapolis, ID).

### S1. The calculation of the estimated cost of digital data storage using de novo DNA synthesis

Based on our calculations, to store 5 minutes of a 1080p YouTube video stream in commercially acquired DNA costs over 382 million dollars, consumes over 100KWh of energy, takes over 4 days, and produces over 15 liters of toxic waste.

The rough size of Youtube HD videos with a resolution of 1080p, based on the video file-size calculator http://toolstudio.io, is 160 MB. On another side, Integrated DNA Technologies (IDT ) reported the cost of synthesizing one base pair of oligo as $0.011.

This information provided the possible cost calculation of storing this file in DNA memory using de novo DNA synthesis.

$$160 \text{ MB} = 1.34e{+}9 \text{ bits} \tag{SE1}$$

If each nucleotide stores 2 Bits, 1.34e+9 bits could be stored in 671,088,640 bp.

The price of storing 5 minutes of a 1080p YouTube video stream is calculated as follows:

$$671{,}088{,}640 \text{ bp} * \$0.011 = \$382 \text{ million} \tag{SE2}$$

### S2. Mechanism of DMOS writer system

DMOS uses a programmed machinery base-editing tool to write the digital data in the form of sequence edits at the precise locations of DMOS registers. To write the data on DMOS bits, we used a DMOS writer system composed of two enzymes: cytidine deaminase mutagenic protein "APOBEC3A" and a mutant form of Cas9 known as dead Cas9 (dCas9). dCas9 is accompanied by a programmed machinery system of gRNA and creates Ribonucleoprotein (RNP). Each DMOS bit consists of a 23bp-long data 'state' section and a unique 40 bp 'index' sequence (Figure. 1).
During the base-editing reaction, the CRISPR effector Cas9 recognizes a 3 bp portion of state defined as a "protospacer adjacent motif, "; resulting in binding to 20bp-long state section within a double-stranded DNA molecule. Next, the 20 bp corresponding gRNA sequence, known as "spacer", recognizes the state section (Fig.1 and S1) and creates a nucleotide "R-loop" structure at the target positions within DMOS bits (Fig.1, S1, S2, S3). The DMOS writer system selectively programs the mutations in the form of sequence edits across the DMOS registers during the writing process. We encode the new data into the existing DNA bits by editing the sequence in the DNA bits through activation or deactivation of the writer system. The writer system uses a cytidine-deaminase that chemically converts
deoxycytosine (dC) to deoxyuracil (dU). Next, dU is converted to deoxythymine (dT) through the active process of writing using Q5U polymerase (Fig.S1). The product of this process is a DNA molecule that contains dT bases instead of dC bases at target regions; as a result, the state of each domain change from intact to mutated. As a further step, we digested the unmodified nucleotide backbone with lambda exonuclease after mutation to increase the likelihood of the modified register being amplified. After exonuclease digestion, we used Q5U polymerase to amplify the remaining modified register. We selected Q5U polymerase to amplify our modified registers, reading the uracil-containing registers in nanopore sequencing reads.
DMOS registers amplified with a reverse 5'-phosphorylated primer, that showed with the red dot in Fig. S1. This modification enables the lambda exonuclease to degrade the unmodified strand of our register, leaving the encoded strand as a register for PCR amplification. Finally, we observe a significant presence of dT in each modified register, indicating successful mutations in our selected bits with no footprint of cross-reactivity (Figure. S1).

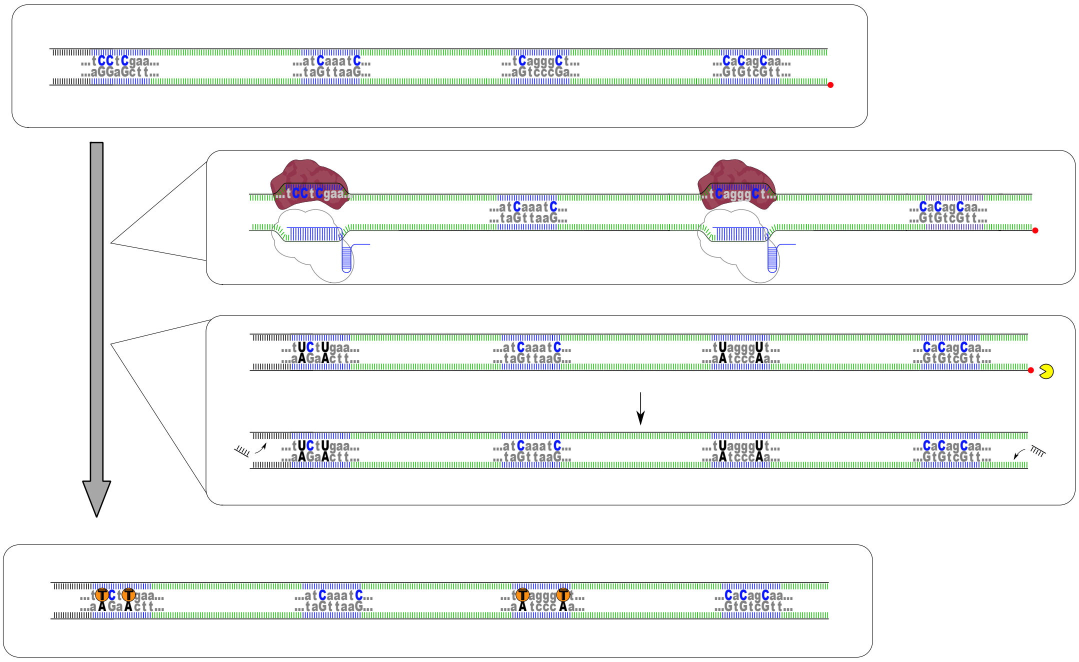

Figure S1. DMOS enzyme encoding protocol. A combination of RNPs target the DMOS register at the desired positions to write the data via a programmable APOBEC3A-driven mutation reaction. The modified register addressed to lambda exonuclease digestion (yellow) and amplified in the active process of writing using Q5U polymerase, and read the uracil nucleotides as thymines in the final sequencing read.

### S3. Cleavage efficiency of gRNA to DMOS bits

To visualize the cleavage efficiency of the spacer sequences of gRNA, we used the active Cas9 proteins and confirmed the targeting of the writer system at the precise location of their complementary domains (Fig. S2). Unlike the dCas9 proteins used in our writing system, the active Cas9 proteins will cleave the DMOS tape at complementary regions. Each digestion splits the tape into two fragments, indicating that the spacer sequences of gRNA show high binding efficiency to the respective state. Therefore, it demonstrates that the gRNA will accord the bits to the R-loop conformation required for APOBEC3A to recognize and deaminate the nucleotides within the loop. We validated the results of digested tape fragments on 2% agarose gel to determine the length of the fragments compared to the control 1108 bp template labeled "C" in the Fig. S2. As shown in Fig. S2, we observed bands of a smaller length relative to the negative DMOS register control. This result indicates that digestion only occurs in precise locations as a function of Cas9. We also observed an "X" pattern across all digestion sites consistent with where we expect the Cas9 protein to cut, showing likely no significant off-target cleavage across the register. This result indicates that the targeting of bits occurs only at the precise location of interest.

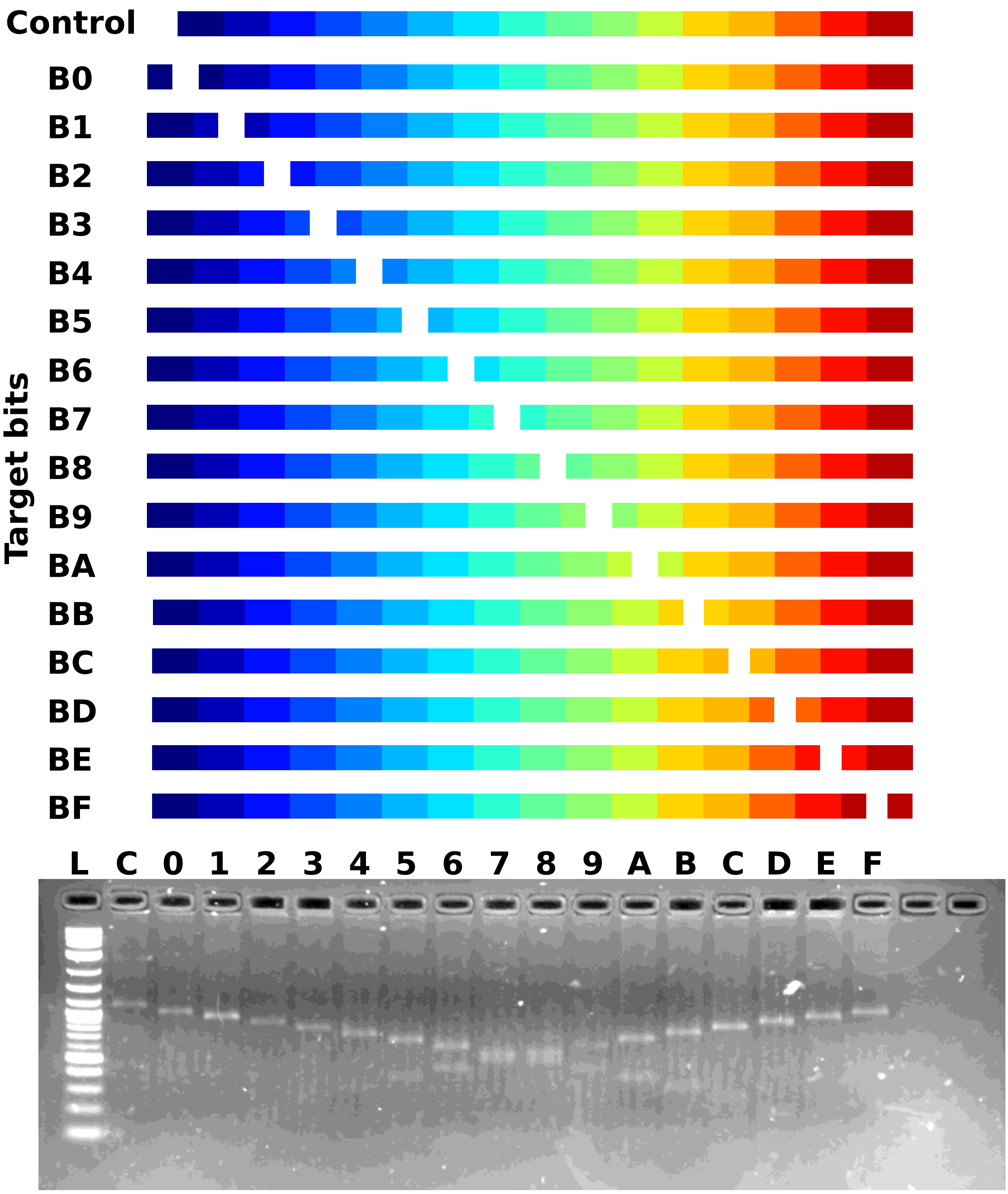

Figure S2: Schematic of efficiency of gRNA to DMOS bits. The efficacy of the writer system was evaluated in the form of Cas9 cleavage activity at the desired bit positions. The gel results confirm that there is no evidence of off-target cleavage, showing the high efficiency of dCas9 for our writing system.

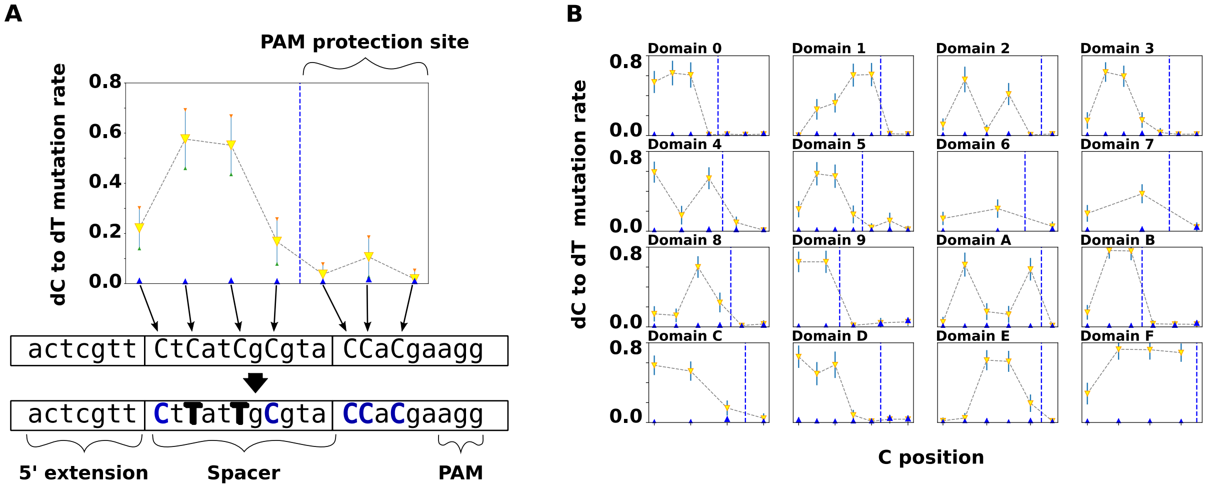

Figure S3: . dC to dT orthogonality analysis: A) Schematic view of mutational signature for one DMOS bit. A mutational signature is depicted as conversion dC's to dT's across the entire 'state' section of a DMOS bit during the base editing reaction. The evidence of mutation appears through changing the state of the bit from '0' (unmutated) to '1' (mutated). The DMOS algorithm deterministically programs the bits and stores the data in the domain states by writing the data on the DMOS bits and making mutations more efficient. To improve the design of DMOS domains, iterative trinucleotides, indicated here as TCA or TCG, were placed at least six nucleotides upstream of the PAM site. B) In vitro APOBEC3A-driven mutational signature of dCs positions tracked across entire DMOS bits (16 domains of DNA register) as a function of dC-to-dT mutation rate (%). Conversion rates showed significant increases up to 80% when the corresponding dCas9 was utilized. Dash line indicated the PAM protection site and number of Cs located in this region.

### S4. Designing address schemes of DMOS registers.

To produce a set of differentiable DMOS registers (Fig. S4A), we initially used the DMOS addressing model as a "Barcode-based addressing scheme" using 36 random barcodes that identify one register from another in the pool (Fig. S4B). In this system, the number of barcodes informs the maximum capacity. To increase data storage capacity, we defined a unique shuffling (permutation) addressing model for the domain position orders (bits) in each individual DMOS register (Fig.S4B), allowing the generation of up to $2.09 \times 10^{13}$ different DMOS registers with high-entropy of permutations (Figure. S4B). The order of DMOS bits determines the "DMOS register address" map. DMOS bits permute independently in random order across the register to identify one register from another in the pool. We refer to a collection of DMOS registers in the pool as "DMOS block," or "DMOS tape" in this study, which contains a library of 48 DMOS registers. (Figure. S4A).

Thus, we defined a lexicographic representation of the permutation order (Fig. S4B), where we map the domain positions as B0 = 0, B1 = 1, …, B15=F. For instance, DMOS register addresses of 0, 1, and 2 represent 0123456789ABCDEF, 0123456789ABCED, and 0123456789ABDFCE, respectively. The maximum theoretical number of DMOS register combinations in one single tube is ~2.09 x10^13, corresponding to the upper limit for addressing locations. Therefore, the permutation addressing scheme enhanced the DMOS system's capacity. According to this scheme, the registers were shuffled only in the order of the last six domains. The issue with this approach is the minimum possibility of variations (low-distance) between the order of consecutive register addresses. The low-distance of this addressing scheme leads to miss-addressing and increases the computation cost (Fig. S4B).

To decrease computational cost during decoding, we suggest the High-distance addressing scheme with the lexicographic representation. This method permutates the order of bit positions in the entire register (Figure. S4B). The High-distance permutation addressing scheme ensures consecutive addresses in the DMOS block with insignificant correlation. Herein, we designed a library of DMOS registers in which the order of domains shuffled across entire registers. For example, the register addresses 0, 1, and 2 represent the combinations such as 915760E3CB2D84AF, 6A51BF32EC074D98, and BC85F6E3072A4D19, respectively. We hypothesized that the High-distance addressing scheme reduces the noise and increases the confidence in our measurements (Figure. S4B).

In this study, we defined one DMOS block as a blank drive containing 48 DMOS registers, in which 32 registers use the High-distance permutation addressing scheme, and the last 16 registers use the Low-distance permutation addressing scheme (Fig. S4A).

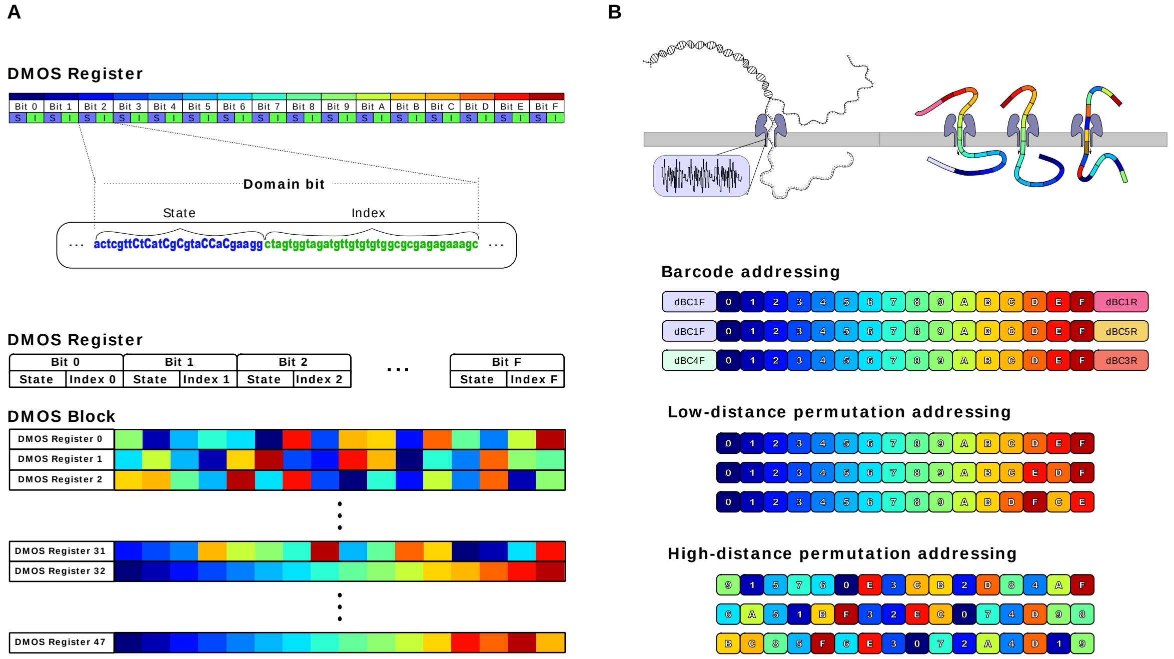

Figure S4: DMOS blank DNA tape. DMOS register defines as a set of 16 predetermined domain bits. As demonstrated in a closer look, a single bit defines as the 'state' section with 23 bp-length and a 40 bp 'index' sequence. To increase data storage capacity, we defined a unique shuffling (permutation) addressing model for the domain position orders (bits) in each DMOS register that generates 48 different combinations of DMOS register as we called "DMOS data blocks" or a "DMOS tape". B) Designing address schemes of the DMOS register. Demonstration of DMOS registers from a trace pool once passing through the nanopore's pores that differentiate three different addressing schemes, including; the Barcoding scheme, Low-distance permutation, and High-distance permutation addressing scheme.

S5. The orthogonality of the DMOS writer system

We introduce a DMOS tape, which uses a programmable, independent, and orthogonal writer system to encode digital data in sequence edits, enabling higher density than the conventional methods. We induced mutations at each bit to analyze the orthogonality and distribution of dC to dT conversions within each bit. We replicated this experiment across the registers, containing 32 different mutation reactions. We observed significant dC to dT conversion rates in multiple dC positions within the written bits. Each DMOS tape in our trials is addressed with barcodes to distinguish registers within the pool and nanopore sequencing run. We collected the dC and dT population data to compare the conversions across the registers from our sequencing (Figure. S4A).

Figure.S3 shows significant dC to dT conversion rates in multiple dC positions within the affected spacers. We also confirmed that dC nucleotides within 6 bp of the protospacer adjacent motif showed an insignificant conversion rate compared to the negative control (Figure.3A). However, the DMOS uses a domain-based coding system that enables information storage in DNA domains rather than individual bases. Thus, we reduce the number of bit errors that may missed in the PAM protection sites within the DMOS bits. This fact emphasizes the robustness of the methods. It bypasses the controversial issues of the current DNA-based memory error correction methods, excluding any bit strings containing insertions, deletions, and degraded information, then performing majority voting to recover the data.[6] The orthogonality experiments have shown efficient conversion of dC to dT within the bits where we applied the corresponding dCas9 complex versus our negative control group. This result also demonstrated no off-target activity in any dCs outside our affected bits, confirming that our encoding strategy is independent, accurate, and precise in encoding digital information into the respective registers (Figure. S5A and B).

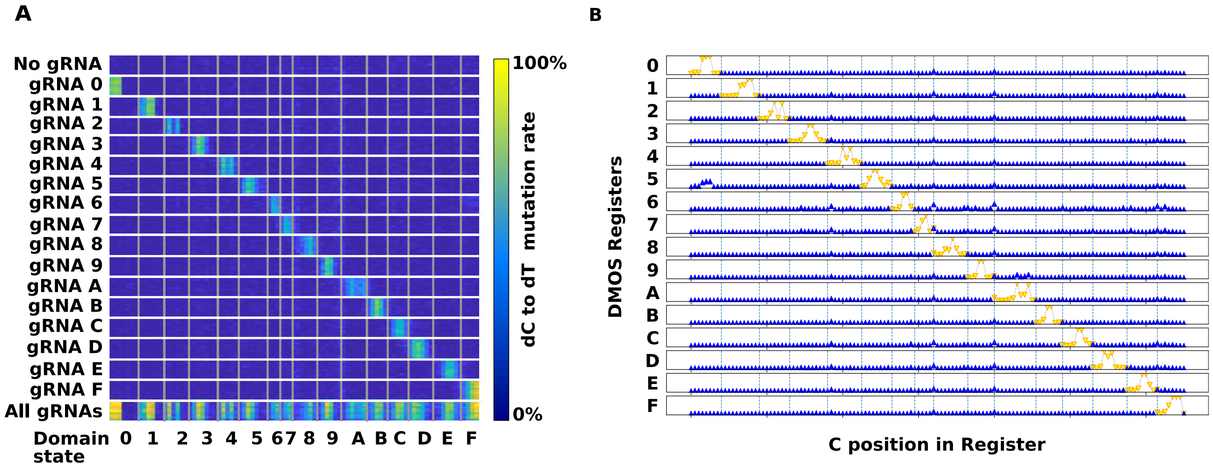

Figure S5: Nanopore sequencing records the orthogonality of DMOS writer system across the domains of DMOS registers.
A) Position-independency of dC to dT ratio determines the modularity of DMOS system to locate the edits at the precise locations across the DMOS registers with no footprint of cross-reactivity. B) Orthogonality analysis of DMOS writer enzyme to assign the cross-reactivity of APOBEC3A/CRISPR-dCas9 across all cytosines as a function of the neighbor bits.

### S6. The called bits

We used a classification model to identify the DMOS register addresses and then mutational states accordingly (Figure. 6). The decoder was developed and written in C++. We first use the Smith-Waterman align algorithm to perform the alignment.[50] Next, the decoder takes the FASTA sequences of the domains B0, B1, …, B15 and determines (pn,sn, cigar) = swalign($B_n$, Seq). Where "p" is defined as positions, "s" as score, "n" as the number of positions [0,1,2,...,15], "$B_n$" is the nucleotide sequence of the n-th domain, and "Seq" is a single nucleotide sequence obtained from the nanopore. Thus, the algorithm generates a list of positions; [p0, p1, …, p15] and sorts them to determine the order of the domains in the DMOS register. Moreover, the decoder uses the Compact Idiosyncratic Gapped Alignment Report (CIGAR report) to generate a string of nucleotides aligned with the cytosines of the bit's states in the FASTA reference. Using the maximum likelihood, the decoder determined each domain's position and alignment score on the sequenced strand. Given the bit order, the decoder generates the lexicographic representation of the address and converts it to an integer number. A valid address number corresponds to our DMOS register's the address library. Whether the address is valid, the decoder determines the values of the mutational states. The DMOS decoder then determines the mutation state. Our DMOS decoder employs two Bayesian steps in a classification decision tree to accomplish this model.

The schematic view of our decoder is shown in Figure.6, demonstrating how the first Bayesian classifier counts all Ts and Cs in the nucleotide sequence of the domain's states. We then accumulate these values for all reads of the same states. Finally, the values used to calculate the dC to dT ratio are as follows:

$$\text{dC to dT} = \text{sum(T)}/(\text{ sum(T)} + \text{sum(C)}) \tag{SE3}$$

We used dC to dT ratio curves to show the states' probability and represent them as unmutated or mutated. We found the regions that curves overlap, creating an uncertain region. Herein, we establish the dC to dT threshold values tu, and tm for unmutated and mutated domain-calls, respectively (Table S3). The classifier determines the state of the mutational state according to the following:

$$\begin{aligned} &\text{Unmutated:} && 0 < \text{dC to dT} < t_u\,, && \text{CS} = -10 \\ &\text{Uncertain:} && t_u < \text{dC to dT} < t_m\,, && \text{CS} = \text{Use second classifier} \\ &\text{Mutated:} && t_m < \text{dC to dT}, && \text{CS} = 10 \end{aligned} \tag{SE4}$$

Where CS is the classification score (CS).

We assessed the called-bits based on the results of the first Bayesian classifier as to whether it determines the state as uncertain. The uncertain calls address further analysis under the next Bayesian classifier. Else, it will assign the CS values as -10 for unmutated and 10 for mutated.

Next, we performed a classification training process for each combination of the domain state's nucleotides according to the expected mutational state. As a result, we generated a library containing the score values per each combination to classify the domain's state as either unmutated or mutated. The second Bayesian classifier receives the scores from the library of the nucleotide string and the CIGAR score.

$$(S_u, S_m)=\text{CheckScores (nucleotide position)} \quad \text{(SE5)}$$

Where Su= unmutated score, Sm= mutated score, and iteratively adds the values to the classification score (CS), as follows:

$$uFraci=(S_u/(S_u+S_m))$$

$$mFraci=(S_m/(S_u+S_m))$$

$$CS=Sum(n=i \text{ to } k) ( \log(mFraci ) - \log (uFraci) )$$

(SE6)

The CS values are in the range [-10, 10]. The Bayesian classifier determines the state of the mutational state according to the following:

| | |
|---|---|
| Unmutated: | CS < 0 |
| Mutated: | CS >= 0 |
| Uncalled: | -5 < CS < 5 |

(SE7)

The decoder uses the CS values to transduce the sequence of nucleotides to the corresponding binary values. The results represent binary 0 and 1 for negative and positive CS values and eliminate the uncalled.

**The correctly-called-bits**

To identify the correctly called bits, we first choose the set of Called bits as determined by the equation:

$$\textit{Called-bits} = 100*((\textit{TotalBits} – \textit{UncalledBits})/\textit{TotalBits}) \%$$

(SE8)

We then utilize a reference file to recognize the errors and subtract them from the set of Called bits:

$$\textit{Correctly-called-bits} = 100*((\textit{TotalBits} – \textit{UncalledBits} – \textit{Errors})/\textit{TotalBits}) \%$$

(SE9)

The aforementioned equations are statistically evaluated using bootstrapping analysis of 250 iterations to produce graphs of Called-bits and correctly-called-bits and the curves were fitted using Origin (as shown in Fig.2 and Fig.3).

Our decoder employs to explore the applications for the tradeoff writing/reading recovery rate and determines whether a long read is required to label a particular state as mutated or unmutated. Hence, we configured our system to take snapshots every 100 intervals, generating the correctly-called-bits analysis as per the reads function in Figures 2 and 3. Depending on whether the DMOS encoder uses error correction, the Python script informs the LDPC decoder algorithm and records the retrieval of stored data. We next add error correction codewords to improve data storage's reliability. We used low-density parity-check (LDPC) codes that set the error threshold close to Shannon capacity limits. The code includes a pre-processing step that adds LDPC error correction to the file's binary contents. [41-44]

The DMOS decoder was written in C++, including a Python interface to generate a real-time visualization of snapshots, and is available on GitHub.(https://github.com/SBMI-LAB/DMOSDecoder).

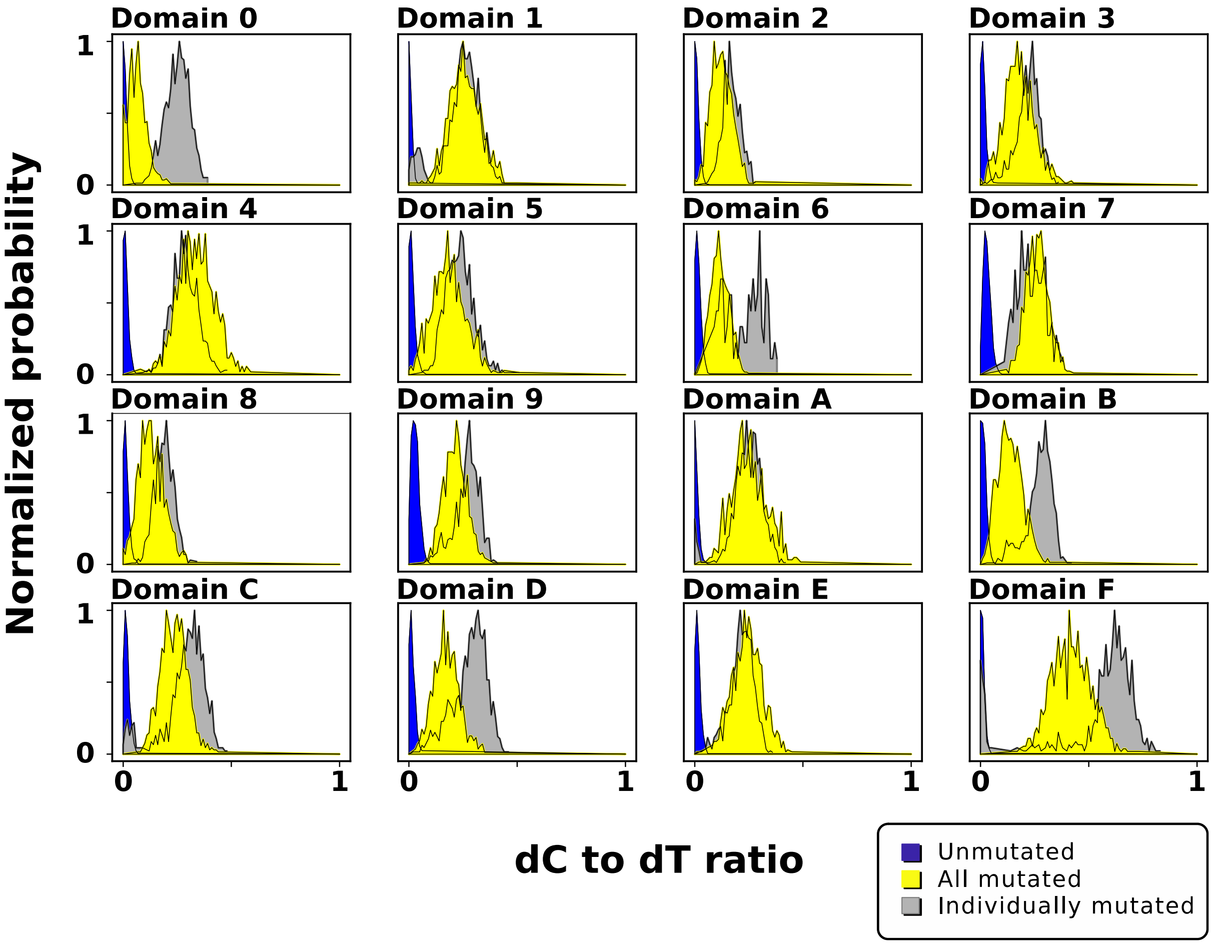

Figure S6: Normalized probability curves of dC to dT ratio to determine the mutation state at the DMOS bit states. Normalized probability distribution of individually mutated (gray) and all domains mutated (yellow) domains compared to unmutated domains (blue).

Curve fitting equation of the called-bits and the correctly-called-bits for Bitmap representation of our school's logo on DMOS tape:

The called bits:

$y = A_1 e^{-reads/t1} + A_2 e^{-reads/t2} + y_0$

$A_1 = -54.7551$
$A_2 = -35.14309$
$t_1 = 625.6624$
$t_2 = 3059.59351$
$y_0 = 92.14322$
$R^2 = 0.99966$

(SE10)

The correctly-called-bits:

$y = A_1 e^{-reads/t1} + A_2 e^{-reads/t2} + y_0$

$A_1 = -57.49084$
$A_2 = -34.94659$
$t_1 = 633.75741$
$t_2 = 3088.54435$
$y_0 = 91.08862$
$R^2 = 0.9996$

(SE11)

Curve fitting equation of the called-bits and the correctly-called-bits for encoding the title of this study on DMOS tape:

The called bits:

$y = A_1 e^{-reads/t1} + A_2 e^{-reads/t2} + y_0$

$A_1 = -54.23359$
$A_2 = -34.61551$
$t_1 = 1373.51658$
$t_2 = 6571.78245$
$y_0 = 89.08068$
$R^2 = 0.999995$

(SE12)

The correctly-called-bits:

$y = A_1 e^{-reads/t1} + A_2 e^{-reads/t2} + y_0$
$A_1 = -55.06187$
$A_2 = -35.42389$
$t_1 = 1447.93899$
$t_2 = 6596.35643$
$y_0 = 86.06278$
$R^2 = 0.99996$

(SE13)

S7. Design of LDPC codes for error protection

We used the Protograph LDPC library (https://github.com/shubhamchandak94/ProtographLDPC)[47][44], to design the LDPC codes for simulation and obtain our error-recovery model subsequently. We designed two types of LDPC codes: Protograph and Regular.

We modified the number of check nodes for the Regular LDPC codes to achieve redundancies ranging from 20% to 90%. For the Protograph LDPC codes, we selected the Protograph type accumulative repeat by 4 jagged accumulate (AR4JA) to define the Generator and Parity Check matrices, with a message-code ratio of 3/4, 2/3, and 1/2 to obtain 25%, 33%, and 50% redundancy, respectively. We noticed that the storage capacity of each LDPC code is inversely proportional to the amount of maximum redundancy overhead added. Our first demonstration of the DMOS writer system encodes 576 bits with 25% redundancy, while it encodes only 77 bits with 90% redundancy (Fig. S8). This result justifies the tradeoff writing/reading to achieve an industrially acceptable data recovery rate.

Protograph LDPC with 25% redundancy uses 576 bits per message (72 bytes) and 768 bits per codeword (96 bytes). We developed a Python script to communicate with the LDPC library that allows the conversion of the intermediate binary files for input/output and records the signals of the LDPC decoder (Fig. S8, S9).

S8. The error-recovery simulation

We ran an error-recovery simulation for different LDPC codes that calculates the decoder's recovery rate per number of errors. The simulator creates a random message according to the capacity of the LDPC codes and creates the codeword. We iteratively inject errors into the codeword and test if the decoder recovers the data. The simulator repeats this procedure 1000 times before increasing the number of errors and recording the chance of recovery per read (Fig.S8).

$$y=A2+(A1-A2)/(1+\exp(Errors-xo)/dx)$$

Where y is Recovery rate (Errors)%
A1 = Initial value of Recovery rate when Errors is 0, A2 = Final value of Recovery rate when Errors is infinity,
Xo = number of Errors where Recovery rate (Xo) approximately reaches 50%, and dx = constant number which determines the slope of the curve close to Xo

*(SE14)*

This simulation creates the error-recovery rate model for different LDPC codes. Given the received errors from the experiments (Fig. 2 and 3), we add reads to a number of errors, feed the model, and represent the DMOS recovery rate (%) per read. Figures 2 and 3 show the error-recovery rate per reading length using the error-rate from the corresponding experiments. Figures S8 show the error-recovery rate per errors % for 1000 iterations by increasing the number of errors and recording the chance of recovery rate % per read. The decoder inform the LDPC codes can recover nearly 100% of the cases. Simulation results imply Protograph LDPC shows a better setting than Regular LDPC with a similar redundancy ratio.

In the Bitmap representation of our School's logo, Protograph LDPC with 25% redundancy showed being capable of recovering the data in nearly 5K reads. We selected this redundancy for further experiments due to the higher number of bits stored in the codeword and assessed the recovery capabilities. Therefore, we encode this study's title using the 25% overhead added. We observed an increased error-rate that required long sequencing reads to achieve a valid recovery, as shown Figure 3. Given this study's error-rate, we simulated the different LDPC codes to determine the amount of required overhead. In good agreement with computational analysis, the experimental result confirmed that using Protograph LDPC with 25% redundancy, we recover the data after 100K (Fig.3), with a 67% probability (Fig.S8). This result justifies the tradeoff between redundancy and data capacity. The simulation study offers the Protograph LDPC code with 50% redundancy and the best error recovery performance (data recovered at 3000 reads) while preserving a decent storage capacity for future works.[41-44]

Curve fitting equation of the bootstrapping error-recovery simulation for Regular LDPC of Bitmap representation of our school's logo on DMOS tape:

$$y=y0+A1* Read^n/K^n+ Read^n \tag{SE15}$$

*Where y0 is statrt point of graph, and A1 is defined as End point of graph and K and N were used as a constant number for curve fitting:*

Curve fitting equation of the bootstrapping error-recovery simulation for Protograph LDPC of our school's logo on DMOS tape: $y=y0+A1* Read^n/K^n+ Read^n$

(SE16)

Curve fitting equation of the bootstrapping error-recovery simulation for Regular LDPC of encoding the title of this study on DMOS tape:

$$y=y0+A1* Read^n/K^n+ Read^n \tag{SE17}$$

Curve fitting equation of the bootstrapping error-recovery simulation for Protograph LDPC of encoding the title of this study on DMOS tape:

$$y=y0+A1* Read^n/K^n+ Read^n \tag{SE18}$$

Table_SE14:

| Redundancy | Value |
|---|---|
| Protograph_25 | A1=99.97827<br>A2=-0.04871<br>x0=24.90464<br>dx=1.47296 |
| Protograph_33 | A1=99.92182<br>A2=0.00564<br>x0=37.87174<br>dx=1.77631 |
| Protograph_50 | A1=100.11285<br>A2=0.38687<br>x0=71.96793<br>dx=2.69267 |
| Regular_30 | A1=100.24655<br>A2=0.0452<br>x0=22.34926<br>dx=1.14607 |
| Regular_40 | A1=100.26255<br>A2=0.04968<br>x0=31.50981<br>dx=1.50235 |
| Regular_50 | A1=100.2023<br>A2=0.09014<br>x0=40.64873<br>dx=1.81469 |
| Regular_60 | A1=100.22736<br>A2=0.11037<br>x0=49.75211<br>dx=2.14945 |
| Regular_70 | A1=100.23028<br>A2=0.08922<br>x0=59.19088<br>dx=2.53653 |
| Regular_80 | A1=100.24714<br>A2=0.2042<br>x0=67.85497<br>dx=2.88074 |
| Regular_90 | A1=100.24409<br>A2=0.73111<br>x0=75.91096<br>dx=2.88074 |

Table_SE15:

| *JSNN_Regular* | *20%* | *30%* | *40%* | *50%* | *60%* | *70%* | *80%* | *90%* |
|---|---|---|---|---|---|---|---|---|
| Start | -2,8145 | -1,19846 | -0,76932 | -0,04302 | -0,81777 | 5,91448 | 31,47089 | 64,27884 |
| End | 95,32917 | 99,92512 | 100,02251 | 100,01691 | 100,01522 | 100,00066 | 100,00003 | 100 |
| K | 3452,38662 | 916,44743 | 453,96099 | 295,81589 | 212,99211 | 172,03107 | 160,49862 | 157,49104 |
| N | 1,72727 | 3,46459 | 4,59958 | 5,60729 | 5,56724 | 7,79442 | 10,70578 | 15,80997 |

Table_SE16:

| *JSNN logo_Protograph* | *25%* | *33%* | *50%* |
|---|---|---|---|
| Start | -1,33597 | -0,59048 | 41,5928 |
| End | 99,8705 | 99,93745 | 99,99796 |
| K | 727,81901 | 329,88765 | 130,51987 |
| N | 3,72265 | 5,25478 | 7,39503 |

Table_SE17:

| *Title of this study_Regular* | *20%* | *30%* | *40%* | *50%* | *60%* | *70%* | *80%* | *90%* |
|---|---|---|---|---|---|---|---|---|
| Start | 0 | -0,10027 | -1,85494 | -0,59461 | -0,28044 | -0,3157 | -0,40654 | -0,34348 |
| End | 0 | 5,48915 | 87,39731 | 998,81142 | 100,00549 | 100,06175 | 100,07247 | 100,02875 |
| K | 0 | 31995,6953 | 17447,2682 | 7827,75856 | 5080,60356 | 3659,35913 | 2859,0301 | 2325,03097 |
| N | 0 | 2,03998 | 2,22827 | 3,98423 | 4,8398 | 5,22339 | 5,4843 | 5,9277 |

Table_SE18:

| *Title of this study_protograph* | *25%* | *33%* | *50%* |
|---|---|---|---|
| Start | -0,4704 | -0,88808 | -0,42944 |
| End | 23,94389 | 98,38785 | 100,01967 |
| K | 33686,3999 | 9363,27747 | 2566,41644 |
| N | 1,84031 | 3,5108 | 5,73664 |

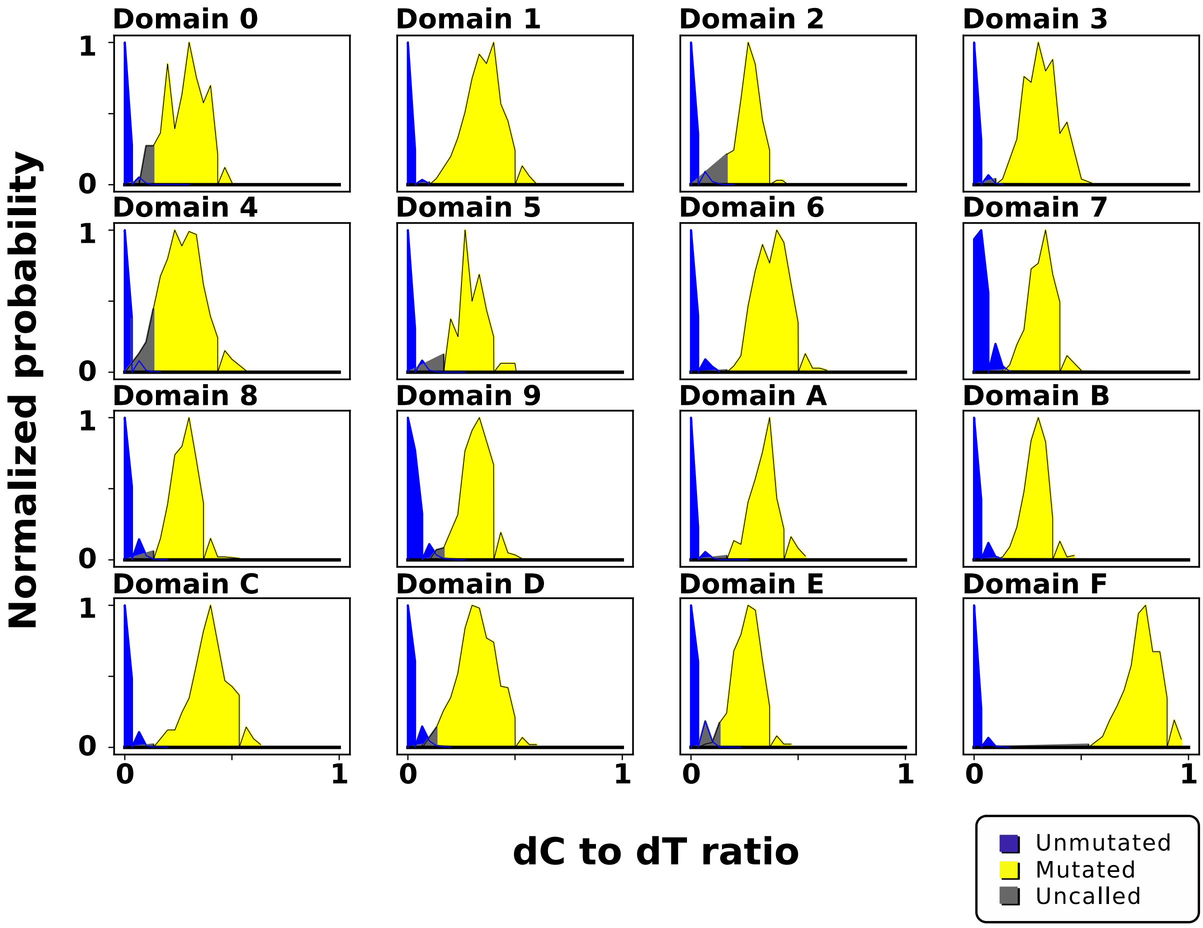

Figure. S7. Normalized probability curves of dC to dT ratio for each individual domain of DMOS registers at bitmap representation of our school's logo. Interception of the intact and mutated curves shows uncertainty in the classification decision tree analysis (threshold shows as gray), and the limits of the interception allow us to determine the threshold values using the first Bayesian classifier.

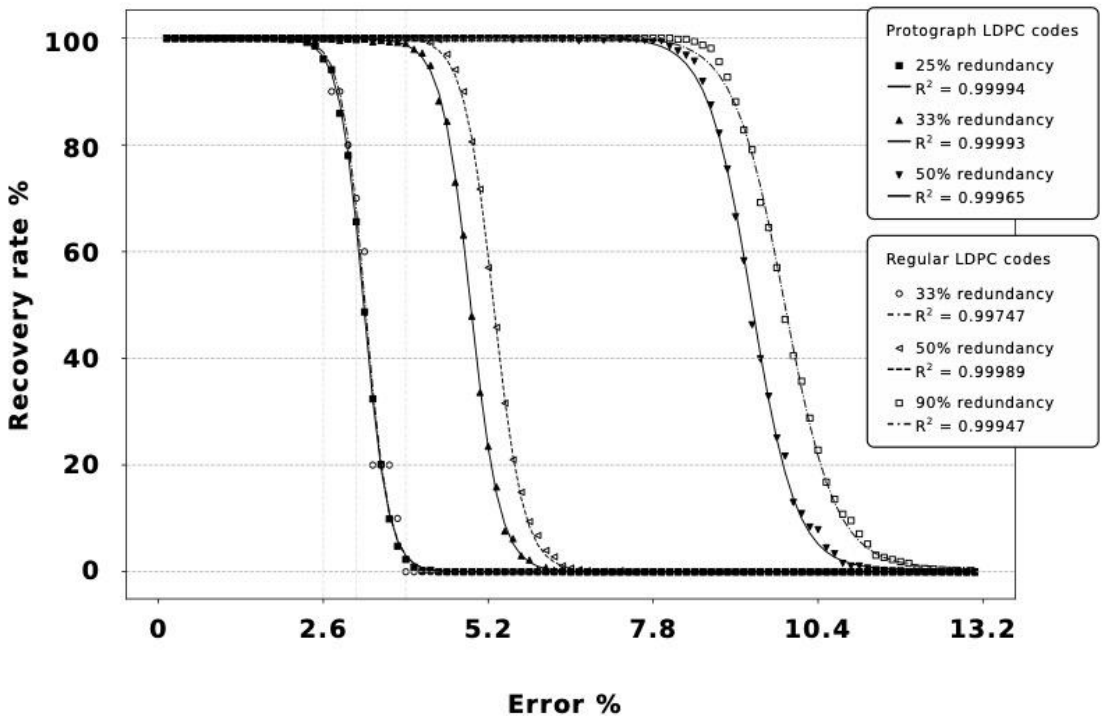

Figure. S8. DMOS error-recovery simulation model for different LDPC codes. The decoder calculates the recovery rate per number of errors for 1000 iterations by increasing the number of errors and recording the chance of recovery rate % per read.

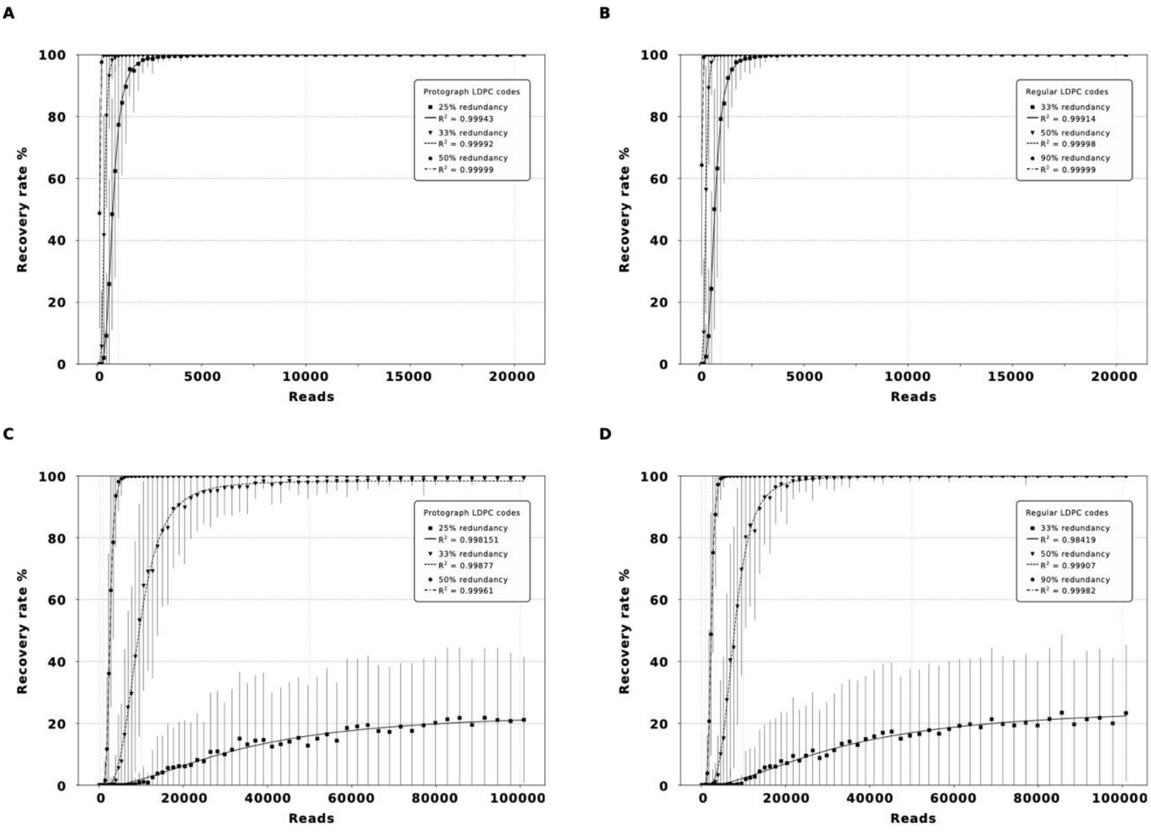

Figure S9. DMOS error-recovery simulation per reads. Error-recovery simulation for different LDPC codes using 1000 repetitions to obtain an error-recovery rate model for each decoder. Data recovery simulation for A) JSNN_Protograph, B) JSNN_Regular, C) Title of this study_Protograph, D)Title of this study_Regular. 3D representation of sequencing reads vs. Protograph LDPC for 33%, and 50% redundancy in the codeword per reads as a function of recovery rate (%). For the Protograph LDPC codes, we used a message-code ratio of 3/4, 2/3, and 1/2 to obtain 25%, 33%, and 50% redundancy, respectively, and 3D representation of sequencing reads vs. Regular LDPC redundancy ratio (%) for 33%-90% redundancy in the codeword as a function of recovery rate (%).

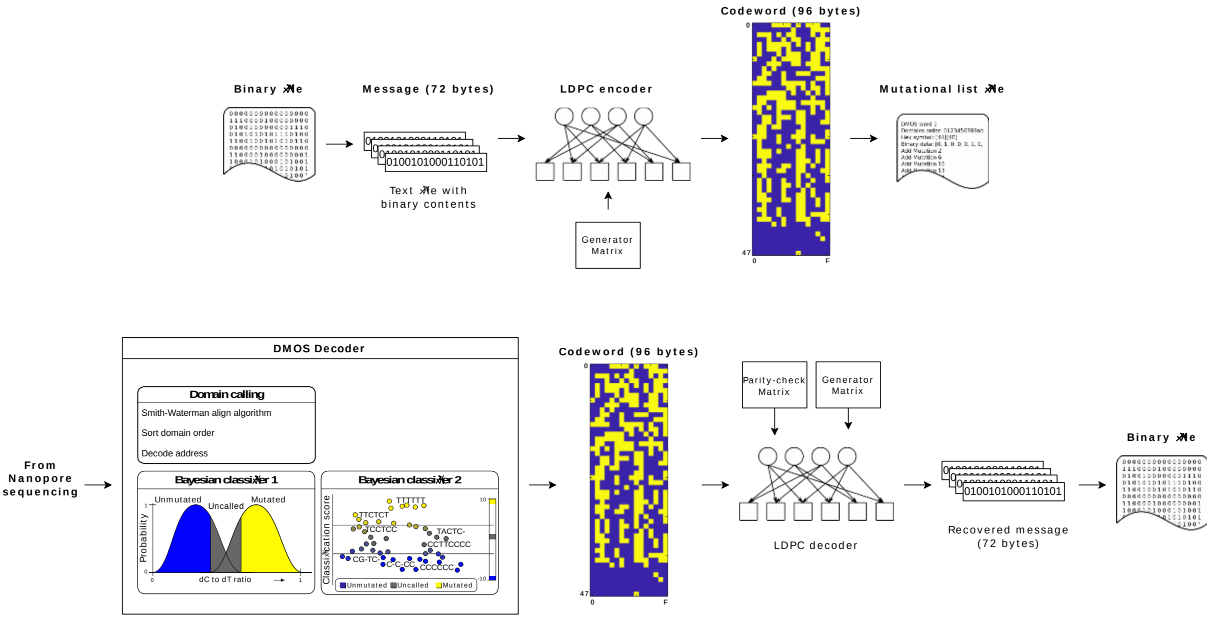

Figure. S10. DMOS coding and encoding  scheme. The DMOS encoder uses binary files and converts it to the byte arrays with codewords. The DMOS encoder then depends on including error correction, transduces the coding protocol to DMOS writer, and change the state of domains from 0 to 1 on DNA tape.

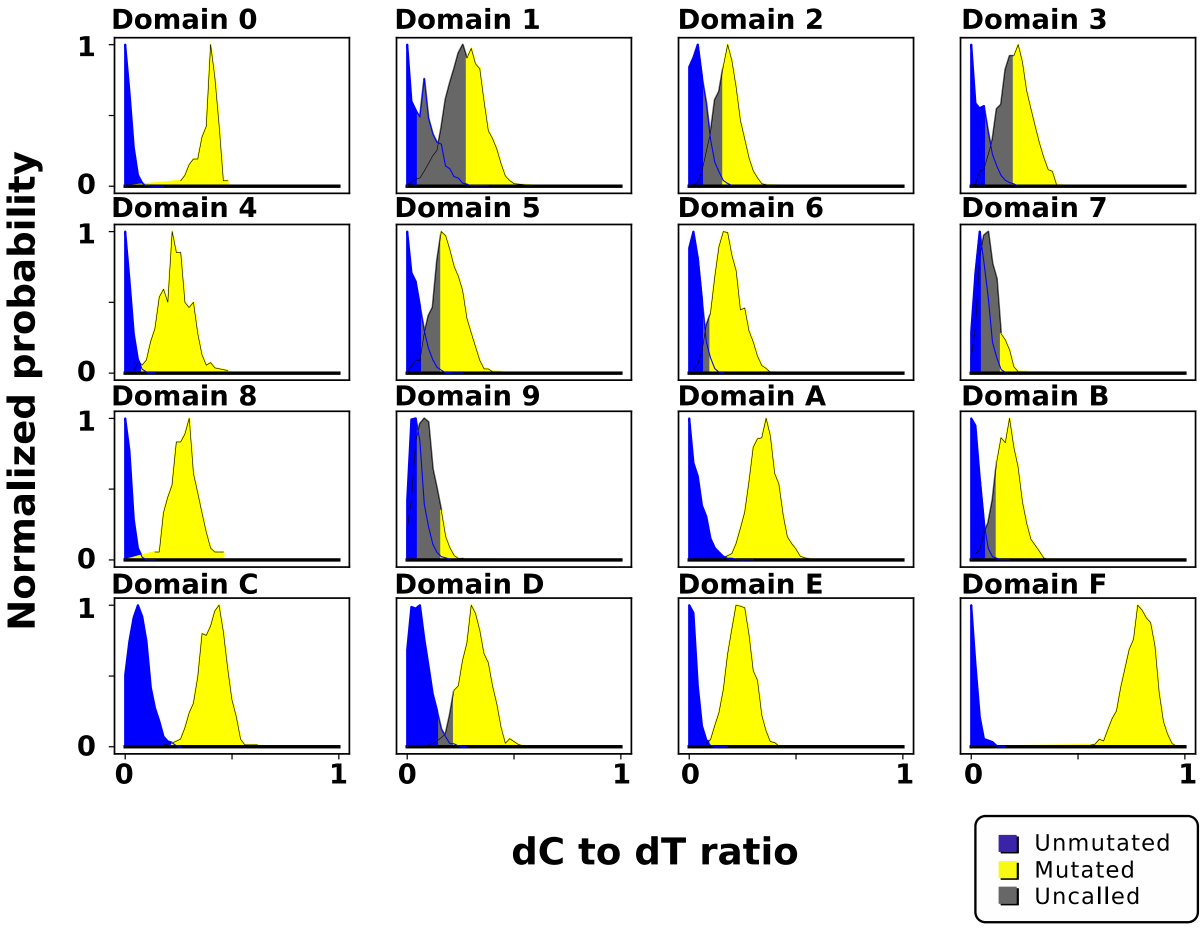

Figure. S11. Normalized probability curves of dC-to-dT ratio for each bit of DMOS registers at the experiment of encoding the title of this study.

**Table S1: The sequences of DMOS bits and indexes**

| | |
|---|---|
| **Initiator** | atcacgaggccctttcgtcttcaagaattc |
| **Index** | TTTATAGAAAACGTTTTGAAGAAGAAGATGATCTCT |
| **State 0** | ACTCactagtcctcgaaaacctcgtgg |
| **Index 0** | TGTCCTACTATGTCTTCTCTCTTCTACTACTTACCT |
| **State 1** | ACTCctccaatcaaatcagtcactagg |
| **Index 1** | GGATGGATGATCCCACACCTCACACGCAGGAGAGAA |
| **State 2** | ACTCtctggtcagggctcggacactgg |
| **Index 2** | CTAGTGGTAGATGTTGTGTGTGGCGCGAGAGAAAGC |
| **State 3** | ACTCtcattcacagcaactgcagcagg |
| **Index 3** | TTGCGACGATGACTGACGACTGCACGAAAAGCTGGA |
| **State 4** | ACTCatggtcaactcaatccaaaatgg |
| **Index 4** | GTGAGGAGGAGAAGTAAAAGAAAGCTTCGAGAGAGT |
| **State 5** | ACTCgttctcatcgcgtaccacgaagg |
| **Index 5** | AGTTTACACGGCGCTCTTTCCGGTTTGATCTTGCAC |
| **State 6** | ACTCatcaatagtgtcatggcatgtgg |
| **Index 6** | ATGTTTACGCACGCGTTTTCCCACCCACGATGTTGT |
| **State 7** | ACTCtcgggagaaaggtcgctgtgagg |
| **Index 7** | CTGTTTGCACACACACCCGCACACCCTGTTCCCTCG |
| **State 8** | ACTCatcacgagttcacgataccgtgg |
| **Index 8** | ATGCGTTGCGTTGTTTTGCGTTCCACACCACACGTT |
| **State 9** | ACTCttgtggtcaatgtcactccgagg |
| **Index 9** | ATCCAAAGAGAACTGGGATTTCTAAAAGAGAGAGAA |
| **State A** | ACTCaagctcagcctcgttaaacgtgg |
| **Index A** | GTTTTACCTTTTGCGCTTTTGTCTTCGTTCGTCCCT |
| **State B** | ACTCgaacagatcatcaacccattagg |
| **Index B** | GGCTCCCTACCACACACCACGTTTTGATGATAGTTG |
| **State C** | ACTCattcaatcaagctgcaaaggtgg |
| **Index C** | TACGAGAGGAAGCTTCACACACCACCACGATCGGAT |
| **State D** | ACTCgattcgaatatctctcttcgagg |
| **Index D** | CTTGCGCACACCTCACACACGTGTTTGTGTTGTGTT |
| **State E** | ACTCgcctcatcagcagaacaagttgg |
| **Index E** | CGATCCGCACACGCACGTCACACCTATCTTACGTGT |
| **State F** | ACTCtcattccagtcaatgtggaaagg |
| **Index F** | GAAGAAAAGAAAGAGAAGAGAAAACTCAAAAGATGA |
| **Terminator** | ACTCatcgataagctttaatgcggtagtttatca |

**Table S2: The sequences of barcode primers**

| | |
|---|---|
| **dBC1F** | TTTCTGTTGGTGCTGATATTGCGTTGTCGGTGTCTTTGTGATCACGAGGCCCTTTCG |
| **dBC2F** | TTTCTGTTGGTGCTGATATTGCCCGTTTGTAGTCGTCTGTATCACGAGGCCCTTTCG |
| **dBC3F** | TTTCTGTTGGTGCTGATATTGCTGTGTCCCAGTTACCAGGATCACGAGGCCCTTTCG |
| **dBC4F** | TTTCTGTTGGTGCTGATATTGCTTCTATCGTGTTTCCCTAATCACGAGGCCCTTTCG |
| **dBC5F** | TTTCTGTTGGTGCTGATATTGCCAGGGTTTGTGTAACCTTATCACGAGGCCCTTTCG |
| **dBC6F** | TTTCTGTTGGTGCTGATATTGCGAACAAACCAAGTTACGTATCACGAGGCCCTTTCG |
| **dBC1R** | TTGCCTGTCGCTCTATCTTCCCGTGGGAATGAATCCTTTGATAAACTACCGCATTAAAGC |
| **dBC2R** | TTGCCTGTCGCTCTATCTTCCAAAGGCAGAAAGTAGTCTGATAAACTACCGCATTAAAGC |
| **dBC3R** | TTGCCTGTCGCTCTATCTTCGCACAGCGAGTCTTGGTTTGATAAACTACCGCATTAAAGC |
| **dBC4R** | TTGCCTGTCGCTCTATCTTCTGAAACCTTTGTCCTCTCTGATAAACTACCGCATTAAAGC |
| **dBC5R** | TTGCCTGTCGCTCTATCTTCTCTATCGGAGGGAATGGATGATAAACTACCGCATTAAAGC |
| **dBC6R** | TTGCCTGTCGCTCTATCTTCGAAAGAAGCAGAATCGGATGATAAACTACCGCATTAAAGC |

***Table S3: The dC to dT threshold values for the first Bayesian classification step**

| Block | Unmutated ($t_u$) | Mutated ($t_m$) |
|---|---|---|
| 0 | 0.077050 | 0.144550 |
| 1 | 0.129758 | 0.178004 |
| 2 | 0.071650 | 0.151862 |
| 3 | 0.113244 | 0.168885 |
| 4 | 0.082900 | 0.166900 |
| 5 | 0.044773 | 0.139677 |
| 6 | 0.070100 | 0.157850 |
| 7 | 0.042900 | 0.094650 |
| 8 | 0.118000 | 0.178000 |
| 9 | 0.024487 | 0.102552 |
| A | 0.092800 | 0.160550 |
| B | 0.080100 | 0.147600 |
| C | 0.080100 | 0.147600 |
| D | 0.068250 | 0.135750 |
| E | 0.092200 | 0.177700 |
| F | 0.192950 | 0.288200 |

*Thresholds determine the state of the domain as Unmutated (dC to dT$<t_u$) and Mutated (dC to dT$> t_m$). The dC to dT values in between ($t_m <$dC to dT$<t_u$) are considered uncertain.
